# Energy Landscape Analysis Reveals Thalamic Modulation of Brain State Transitions During Movie Watching

**DOI:** 10.64898/2025.11.30.691450

**Authors:** Qiuyi Liu, Lili Sun, Xinyi Zhao, Wenbin Qu, Jiaqi Zhou, Ziang Wang, Kaizhou Li, Huiting Lei, Xia Liang

## Abstract

Understanding how large-scale brain networks dynamically coordinate during complex, real-world experiences remains a crucial topic in contemporary neuroscience. Leveraging naturalistic paradigm, this study employed energy landscape modeling to characterize the temporal dynamics of brain state transitions and their functional significance. Using the Sherlock fMRI dataset, we identified two dominant brain states: a perception- and attention-oriented state, and an introspective, integrative state, each associated with distinct canonical brain networks. State transitions were differentially associated with the recruitment of these networks and manifested as either the “easy” or “hard” route, depending on the energy required for the transition. Notably, we demonstrated that the probability of specific state transitions robustly predicted inter-subject correlation within three clusters: subcortical-rich, salience/attention-rich, and DMN-rich regions. This indicates that flexible reorganization of cortical networks underlies shared neural engagement during ecologically valid experiences. Critically, the thalamus emerged as a central modulator, displaying activity changes and dynamic nucleus-specific thalamocortical connectivity changes tied to distinct transition profiles. Together, our findings shed light on the energetic and network mechanisms that orchestrate brain dynamics during naturalistic cognition and underscore the pivotal role of cortico-thalamo-cortical circuits in governing flexible, collective neural responses to richly structured real-life stimuli.

## Introduction

Understanding the dynamic coordination of large-scale brain networks remains a central challenge in neuroscience (Shine and Poldrack, 2018). Naturalistic paradigms, such as movie watching, provide an ecologically valid framework for probing the brain’s intrinsic dynamics as they unfold in real time (Baldassano et al., 2017; Chen et al., 2017; Finn and Bandettini, 2021; Hasson et al., 2015; Kringelbach et al., 2023). Recent advances in computational modeling, such as the energy landscape frameworks, enable the characterization of brain state repertoires and their transitions, which can be conceptualized as traversals between stable states (valleys) and state boundaries (hills) within a multidimensional landscape (Munn et al., 2021; Watanabe et al., 2014). This kind of mathematical approach has elucidated how reconfigurations among distributed cortical networks may underpin cognitive flexibility and disability (Shine et al., 2021; Watanabe, 2021; Watanabe and Rees, 2017). Nevertheless, despite these notable computational achievements, the fundamental question of functional relevance, the relationship between such brain dynamics and coordinated neural processes underlying meaningful, naturalistic experiences, remains inadequately explored.

Recent research has increasingly utilized inter-subject correlation (ISC) or its derivatives, such as inter-subject functional connectivity (ISFC), as measures of shared neural responses among individuals exposed to naturalistic stimuli (Chang et al., 2021; Chen et al., 2020; Di et al., 2022; Finn et al., 2020; Lahnakoski et al., 2014; Parkinson et al., 2018; Yeshurun et al., 2017). Accumulating evidence indicates that regions including the primary/secondary visual/auditory areas and association cortices display robust ISC during movie viewing, reflecting highly aligned temporal dynamics of BOLD activity across participants (Chen et al., 2017; Hasson et al., 2004). These synchronized neural patterns not only support shared comprehension and socioemotional engagement with narratives (Finn et al., 2018; Finn et al., 2020; Jangraw et al., 2023; Yeshurun et al., 2021) but are also predictive of individual behavioral outcomes such as memory efficacy and affective responses (Da Silva Ferreira Barreto et al., 2020; Nastase et al., 2019). Building on this, the present study employs ISC as a functional metric to assess whether, and how, transitions between distinct cortical brain states modulate the degree of shared neural processing under naturalistic conditions. By focusing on the temporal coupling between dynamic brain states and ISC, we sought to clarify whether the flexible reorganization of large-scale cortical networks plays a mechanistic role in facilitating collective neural phenomena in ecologically relevant contexts.

Notably, while cortical dynamics may influence ISC and thus have measurable functional significance, these same dynamics are subject to regulation by subcortical structures such as the thalamus (Buckner and DiNicola, 2019; Redinbaugh et al., 2020; Whyte et al., 2024). The thalamus has emerged as a pivotal hub in modulating large-scale brain network dynamics, integrating computations across distributed cortical modules and coordinating adaptive responses to the changing demands of cognition and arousal (Greene et al., 2020; Muller et al., 2023; Shine, 2021; Shine et al., 2023). Recent empirical studies have demonstrated transient modulation of thalamic activity aligning with rapid cortical activity shifts during cognitive tasks or arousal states (Rikhye et al., 2018; Setzer et al., 2022), highlighting the nuanced interplay between specific thalamic circuits and brain state transitions. At a finer microcircuit level, distinct thalamic neuron populations, principally the PVALB-rich core cells and CALB1-rich matrix cells, differ in their projection patterns and likely in their specialized influence on sensory versus integrative cortical functions (Honjoh et al., 2018; Hwang et al., 2021; Jones, 2001; Müller et al., 2020). Specifically, core cells project discretely to sensory cortices, facilitating the precise relay of sensory information, while matrix cells broadcast more diffusely to integrative association regions, supporting higher-order cognitive processes. Nonetheless, the detailed mechanisms through which specific thalamic circuits interface with dynamic brain state transitions remain an open question ripe for further investigation.

In this study, we leveraged energy landscape methodologies to systematically disentangle the temporal brain dynamics in the *Sherlock* dataset, a rich fMRI resource involving participants engaged with naturalistic audiovisual stimuli, the first 48 minutes of the BBC’s TV series “Sherlock”. By constructing the energy landscape during movie watching, we identified multiple metastable brain states and assessed how the brain transitions among these states over time. We specifically analyzed region-specific neural activity during both “easy” and “hard” state transitions to pinpoint the neural substrates implicated in rapid, large-scale reconfigurations of network architecture in response to dynamic movie stimuli. To elucidate the functional significance of these state transitions, we examined the extent to which specific dynamic brain trajectories predict regional inter-subject correlation. Employing data-driven techniques including LASSO regression and hierarchical clustering, we determined which patterns of brain state transitions most robustly influence ISC distributions and how these effects are organized across canonical brain networks. Finally, recognizing the central modulatory role of the thalamus, we also investigated its unique contribution to brain state dynamics, focusing on thalamic involvement across diverse transition profiles and its functional interplay with major cortical systems. Collectively, these analyses provide new insight into the energetic and network principles underlying brain dynamics in naturalistic settings.

## Materials and Methods

### Sherlock dataset description

This study reports findings from a naturalistic fMRI dataset, the ***Sherlock*** dataset (Chen et al., 2017), which was downloaded from Princeton University’s DataSpace repository (http://arks.princeton.edu/ark:/88435/dsp01nz8062179). The original study included 22 participants (12 males, 10 females) between the ages of 18 and 26, who were native English speakers. Data from 17 participants met the requirements of the original study and were used in this analysis. The naturalistic movie stimulus consisted of the first 48 minutes of the BBC’s TV series “Sherlock”, divided into two parts (about 23 and 25 minutes, 946 TRs and 1030 TRs). MRI was performed on a 3T full-body scanner (Siemens Skyra) with a 20-channel head coil. Functional imaging data were acquired using a T2*-weighted echo planar imaging (EPI) pulse sequence (TR 1500 ms, TE 28 ms, flip angle 64, whole-brain coverage 27 slices of 4 mm thickness, in-plane resolution 3×3 mm^2^, FOV 192×3×192 mm). Anatomical images were acquired using a T1-weighted MPRAGE pulse sequence (0.89 mm^3^ resolution). Standard preprocessing procedures, including slice time correction, motion correction, linear detrending, high-pass filtering (140 s cutoff), and normalization to the MNI template, had already been performed with FSL before downloading. All participants provided informed written consent prior to the start of the study in accordance with experimental procedures approved by the Princeton University Institutional Review Board. Detailed information about this dataset is available in the original report by Chen et al.(Chen et al., 2017).

### Energy Landscape Model Analysis

Before energy landscape construction, we applied the Schaefer (Schaefer et al., 2018) cortical parcellation (100 parcels, 7 canonical networks) to define regions of interest (ROIs). Following previous studies (Ezaki et al., 2017; Watanabe et al., 2014; Watanabe and Rees, 2017), we firstly computed the averaged signal for each ROI. Then, the network-wise averaged signal was computed by averaging the signals of ROIs in each of the 7 canonical networks. Finally, binarization should be performed on the network-wide signal by comparing it to a threshold, typically the mean network activity across the entire scan duration. However, this approach is not ideal for naturalistic datasets, where the global signal is often retained due to its potential physiological significance and the task-related information it may contain (Chen et al., 2017; Simony et al., 2016; Zadbood et al., 2022). To prevent the binarization process from being influenced by this characteristic of naturalistic data, as illustrated in Figure 1A, we adopted an alternative approach. Specifically, at each time point, we compared the network-wise signal (colored lines) to the cross-network mean signal (black line). The binary activity of network *i* at time *t* was denoted as 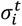, where 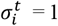 (above average) or -1 (below average). This approach maintained a diverse range of binarized cortical states in data where the global signal was preserved. The energy landscape was estimated by a pair-wise Maximum Entropy Model (MEM), which is fitted using the above processed binary data. The pattern at time *t* was described as 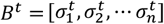, where 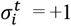 or -1 and represents the binary activity of network *i* at time *t*, and *N* denotes the number of networks (*N* = 7). The empirical probability of a certain pattern *B*_*k*_ (_*k*_ = 1,2, …, 2^*N*^)is constrained by the Boltzmann distribution and given by

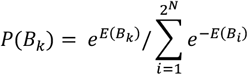

Where E(B_k_) is the energy of activity pattern B_k_, given by

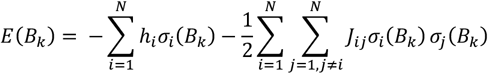

Where *σ*_*i*_(*B*_*k*_)denotes the binarized activity of region *i* when the pattern is *B*_*k*_. Here, *h*_*i*_ and *J*_*iij*_ are the parameter of the model. A gradient ascent algorithm was applied to adjust them, until the model-based mean network activity ⟨*σ*_*i*_⟩_*model*_ and the model-based mean pairwise interaction ⟨*σ*_*i*_*σ*_*j*_⟩_*model*_ were close to empirical observations ⟨*σ*_*i*_⟩_*empirical*_ and ⟨*σ*_*i*_*σ*_*j*_⟩_*empirical*_, respectively (Ezaki et al., 2017; Watanabe and Rees, 2017; Yamashita et al., 2021). In this study, the Sherlock dataset resulted in an energy landscape consisting of 7 local minima which indicate 7 energy basins (128 binarized states were distributed in these basins). For easier understanding, we defined these 7 basins as states 1 to 7 in the rest of our manuscript. The brain configurations of these states were displayed in Figure 1B.

**Figure 1.**
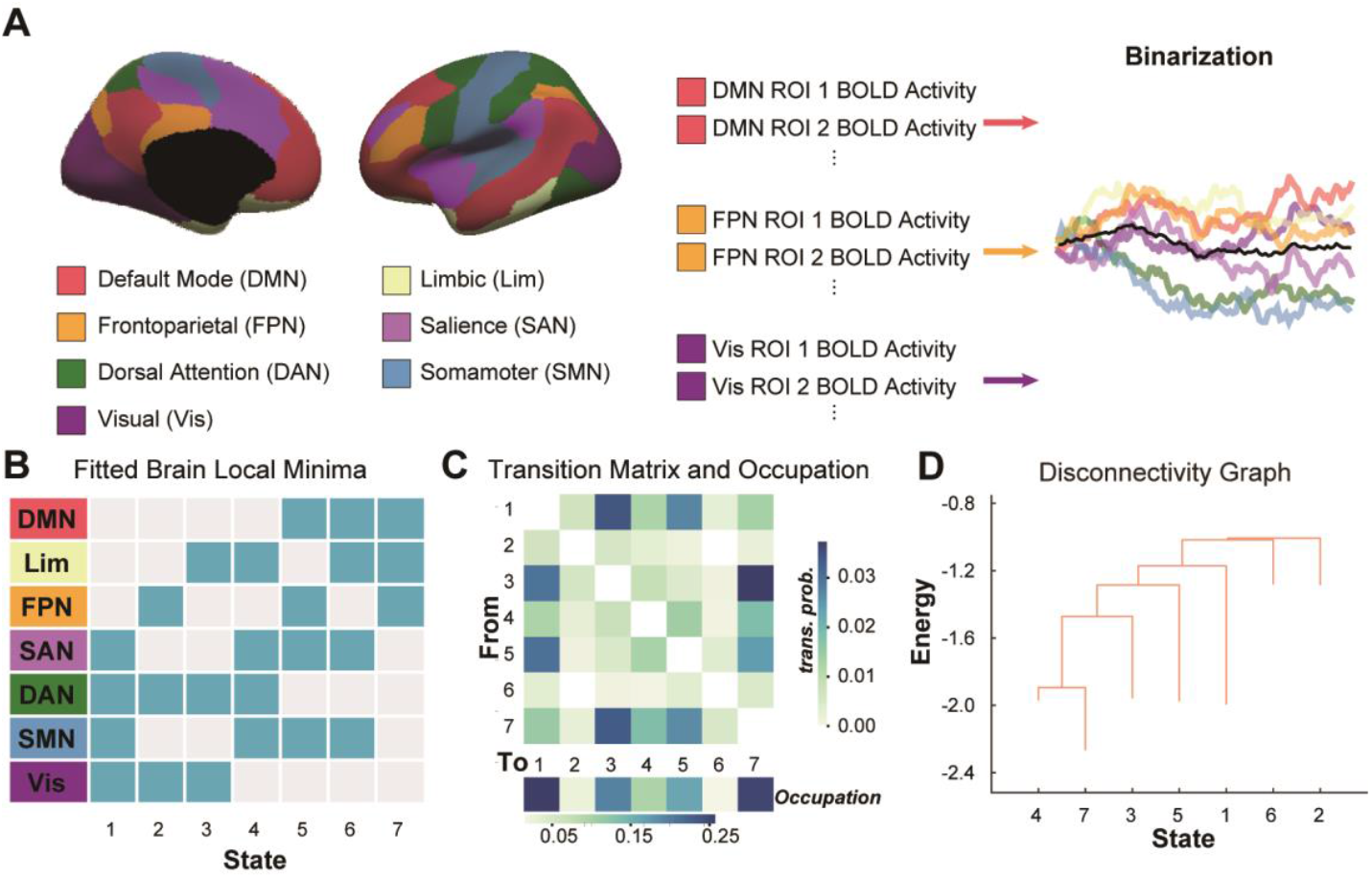
Fitting the Sherlock dataset using the pairwise maximum entropy model. (A) The cortical ROIs were classified into seven functionally distinct brain networks, with network activity binarized using the network-averaged signal. The black line shows the mean activity across all seven networks. (B) Seven brain states were identified using the pairwise maximum entropy model; active networks are highlighted in blue, and each network activation pattern corresponds to a local energy minimum for that state. (C) Group-averaged transition probabilities and state occupancy are shown. Self-state transitions were excluded from the transition matrix and are left uncolored. (D) The relationships among the activity patterns corresponding to energy local minima are illustrated in a disconnectivity graph.

### Energy Landscape Transitions and the Neural Signatures of Hard and Easy Routes

When the brain moves between different states within the energy landscape, these transitions create changes in energy relative to the previous state. Building on a recent study (Luppi et al., 2024), we identified transitions where energy levels rise or fall sharply, labeling these as “Hard” or “Easy” energy routes, respectively. Specifically, energy change was quantified as the difference in calculated energy values between consecutive time points. Transitions with energy changes greater than +2 or less than -2 standard deviations from the individual’s average energy changes were defined as Hard or Easy routes, respectively. Note that the “energy” described in the energy landscape is not biological energy, but rather a statistical transformation of probability distributions derived from the Boltzmann distribution, where low-energy states correspond to high-probability network configurations.

To identify brain regions that potentially support these routes, we analyzed BOLD activity changes around the offset of the Hard and Easy routes, respectively. To characterize BOLD dynamics surrounding these transitions, we used a 21-TR window centered on the route offset, defined as the time point marking completion of a Hard or Easy transition (offset = 0 TR, onset = – 1 TR, the transition offset corresponds to timepoint t, marking the entry into the new state). This longer window was used to stabilize signal normalization and avoid boundary artifacts when z-scoring the data. Within the 21-TR span (–10 TRs to +10 TRs around the offset), BOLD signals of each voxel were z-scored relative to that local window to account for slow drifts.

For statistical comparison, we focused on a 13-TR segment (–6 TRs to +6 TRs around the offset), which captures the canonical hemodynamic response surrounding the transition. Within this segment, the pre-route period was defined as the average BOLD across –6 to –1 TRs (before offset), and the post-route period as the average across +1 to +6 TRs (after offset). For each cortical gray-matter voxel (averaged within a 3 × 3 × 3 voxel neighborhood, step = 1), paired t-tests were performed to compare post-route versus pre-route BOLD activity separately for Hard and Easy routes.

### Inter-subject Correlation

The inter-subject correlation is commonly used to identify brain regions with shared activity patterns across subjects. We computed the averaged signal for each ROI of each subject, then calculated the ROI-wise ISC with an open-source algorithm (Nastase et al., 2019). The ISC values of each subject were calculated with a leave-one-out approach. Specifically, the ISC was calculated by correlating the activity of the leave-out subject and the averaged activity across remained subjects. 100 cortical ROIs (Schaefer et al., 2018) and 14 sub-cortical ROIs (Klein and Tourville, 2012) were used in this study.

### LASSO Regression Analysis

To investigate the relationship between cortical state transition probabilities and ISC, we employed LASSO regression with comprehensive cross-validation. The model is formulated by the following objective function:

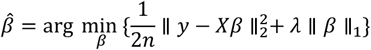

Here, *y*∈ ℝ^*n*^ denotes the vector of ISC values, and *X* ∈ ℝ^*n*×*p*^ is the design matrix composed of *p* transition probability features. The vector *β∈ ℝ*_*p*_ represents the regression coefficients to be estimated, and 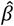 is the resulting estimate obtained through the optimization. The number of subjects is ^*n*^ = 17, and the hyperparameter λcontrols the strength of L1-regularization.

Given the limited sample size (N=17), we employed leave-one-out cross-validation (LOOCV) to prevent overfitting and evaluate feature reliability. For each ROI, the regularization parameter λ was optimized by testing 200 values spaced logarithmically over a range from 10−^5^ to 10 °, selecting the value that minimized the mean squared error. Using the optimal λ identified for each ROI, we constructed 17 LOOCV models, each trained on 16 subjects and tested on the held-out subject. Model performance was assessed using the coefficient of determination:

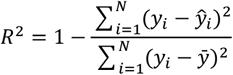

where *yi* represents the actual ISC value for subject *i*, 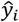denotes the predicted ISC when subject *i* was left out during training, and 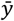 is the mean ISC across all subjects for the given ROI.

To assess statistical significance, we performed 1000 permutation tests. In each permutation, transition probabilities were randomly shuffled at the group level while preserving the full LASSO-LOOCV procedure. ROIs exhibiting R^2^ values significantly higher than the permuted distribution (p < 0.05, FDR-corrected across 114 ROIs) were deemed to be significantly modulated by at least one state transition.

To enhance robustness, we defined stable features as state transition probabilities that were consistently selected (i.e., assigned a non-zero coefficient) across all 17 LOOCV models. This procedure substantially reduced the dimensionality of both ROIs and features. We then applied hierarchical clustering to the resulting matrix of LASSO regression coefficients, where each row represented one significant ROI and each column represented one stable state transition probability. The value in each cell was the coefficient (*β*) quantifying the relationship between that transition and ISC in that ROI. This approach allowed us to test whether ROIs within the same canonical brain networks exhibited similar multivariate profiles of dependence on specific state transitions.

### Thalamic BOLD Activity Dynamics Surrounding Cortical State Transitions

We next investigated alterations in thalamic BOLD activity patterns associated with identified state transitions. To characterize neural dynamics surrounding these transitions, we extracted BOLD signals within a 13-TR window (spanning 6 TRs before to 6 TRs after each transition). The 13-TR window was selected to capture the full hemodynamic response around each transition while minimizing contamination from adjacent transitions of the same type. The extracted BOLD time series for each subject were z-scored and averaged across all transition windows. To establish a statistical baseline at each time point relative to the transition, we conducted 1000 permutation tests by randomly shuffling transition timings, preserving their sequential structure. For each permutation, we computed the mean z-scored thalamic BOLD activity, generating a null distribution of thalamic activity values for each time point. The observed z-scored thalamic BOLD activity at each TR was then compared to its corresponding null distribution using a independent t-test. Significance was defined at p < 0.001/13, with Bonferroni correction for multiple comparisons across the 13 time points.

### Thalamocortical Connectivity Dynamics During State Transitions

To investigate how thalamocortical connectivity changes around state transitions, we computed dynamic functional connectivity (dFC) between two principal thalamic cell populations (Matrix and Core cells) and cortical voxels. The anatomical delineation of these populations was based on a validated template leveraging differential expression of calcium-binding proteins, with Matrix regions identified by high calbindin expression and Core regions by high parvalbumin expression (Müller et al., 2020). fMRI signals for connectivity analysis were extracted from four corresponding thalamic regions: left core, left matrix, right core, and right matrix.

Consistent with our previous analysis, we employed a sliding window of 13 TRs centered on each state transition to compute dFC. This process was repeated for every identified state transition. For each subject, the dFC estimates were averaged across all transition windows. To assess statistical significance, we compared the observed mean connectivity values against a null distribution generated from 1,000 permutations, in which the timing of state transitions was randomly shuffled while preserving the underlying BOLD time series structure. Clusters were defined at P < 0.05 (uncorrected, two-tailed) and tested for significance at P < 0.05 (FWE-corrected, cluster level).

## Results

### Energy Landscape Analysis Reveals Dominant Metastable States in the Sherlock Dataset

We constructed an energy landscape for the Sherlock dataset using an open-source pairwise maximum entropy model (model accuracy: 98.32%, Figure S1A), revealing seven distinct metastable states (States 1-7) that correspond to topographical local minima in the energy landscape (Figure 1B). Among these, State 1 and State 7 emerged as the two most dominant states, accounting for 25.7% and 25.1% of the observed time course, respectively (group-averaged, Figure 1C, bottom). In comparison, States 3, 4, and 5 accounted for 19.4%, 9.5%, and 16.3%, respectively. State 2 and State 6 were visited less frequently, occupying only 2.5% and 1.5% of the temporal dynamics, respectively. Notably, State 1 is primarily characterized by activations in the primary (visual network [Vis] and sensorimotor network [SMN]) and attention networks (dorsal attention network [DAN] and salience network [SAN]), whereas State 7 consistently activates the high-order networks (frontoparietal network [FPN], default mode network [DMN], and limbic network [Lim]). Transitions between State 1 and State 7 were less frequent compared to transitions from either State 1 or State 7 to States 3 or 5 (Figure 1C, top). Meanwhile, the disconnectivity graph (Figure 1D), which describes the energy barrier between any two states, also suggests the transition between State 1 and State 7 would be trapped in other States between them (e.g., State 3, 4, 5).

### Rapid Energy Changes Indicate Certain Regional BOLD Activity Changes

Having identified the brain metastable stases and mapped their associated energy landscapes, we next investigated whether certain brain regions are particularly affected by rapid energy fluctuations during dynamic state transitions. Specifically, we aimed to characterize changes in brain activity along two types of trajectories: “easy routes”, which refer to transitions toward more accessible configurations (i.e., local minima or frequently visited states) that produce decreases in energy, and “hard routes”, which are transitions toward less accessible configurations accompanied by increases in energy (Figure 2A). The analysis revealed that, during easy routes, there was a significant increase in post-transition BOLD activity compared with pre-transition levels in the middle temporal gyrus and the precuneus (FWE corrected P < 0.05; see Figure 2B, top). In contrast, for hard routes, post-transition BOLD activity was significantly higher than pre-transition activity in the left inferior parietal gyrus and the left inferior frontal gyrus (FWE corrected P < 0.05; Figure 2B, bottom).

**Figure 2.**
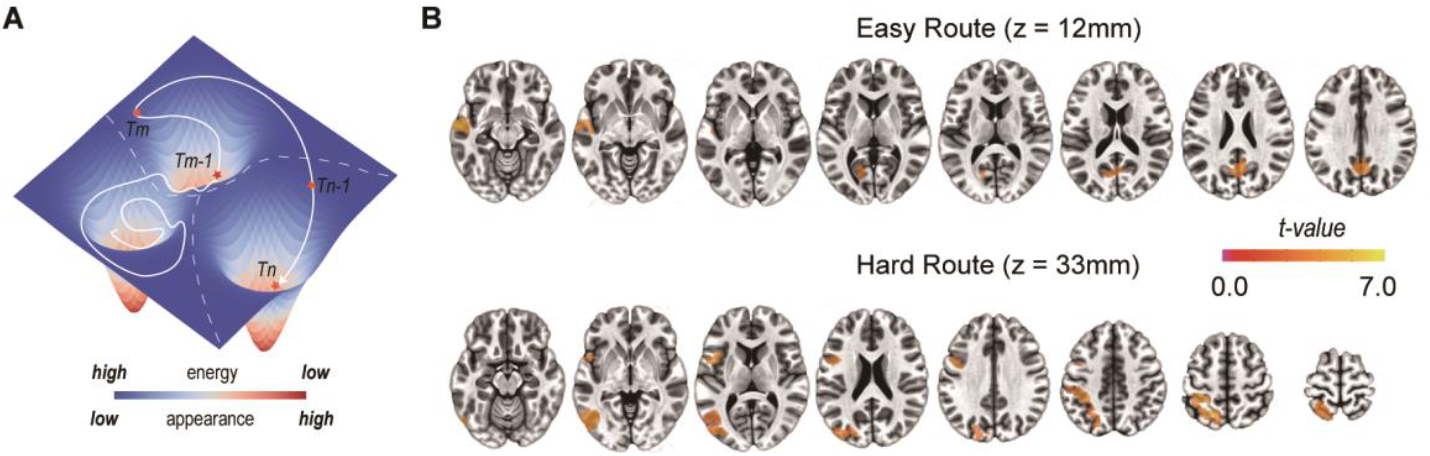
Route-related BOLD activity. (A) Example energy landscape illustrating the route of attractor movement across three local minima. The transition from T_m-1_ to T_m_ represents a hard route, characterized by a rapid increase in energy, while the transition from T_n-1_ to T_n_ represents an easy route, marked by a sharp energy decrease. (B) Clusters demonstrating significantly increased BOLD activity following the transition, as identified by paired t-tests comparing post- and pre-transition BOLD values.

### Relationship Between Brain Dynamics and ISC

To further elucidate the functional significance of the dynamic brain state transitions, we turned our attention to patterns of intersubject correlation. This allowed us to explore how intersubject synchronized neural dynamics might relate to the observed energy fluctuations associated with brain state transitions. When examining ISC values across 114 ROIs as depicted in Figure 3A, we observed particularly strong correlations in visual areas, temporal and medial parietal regions, consistent with previous studies (Chen et al., 2017; Hasson et al., 2004). To investigate how brain dynamics might influence these synchronization patterns, we employed LASSO regression to assess how specific brain state transitions predict regional ISC. As shown in Figure 3B, LASSO regression analysis demonstrated that brain state transitions significantly predicted ISC values in 29 ROIs (P < 0.05, FDR-corrected within 114 ROIs). The predictive performance varied across regions, with R^2^ values ranging from 0.22 to 0.69, indicating moderate to strong relationships between specific transition patterns and ISC.

**Figure 3.**
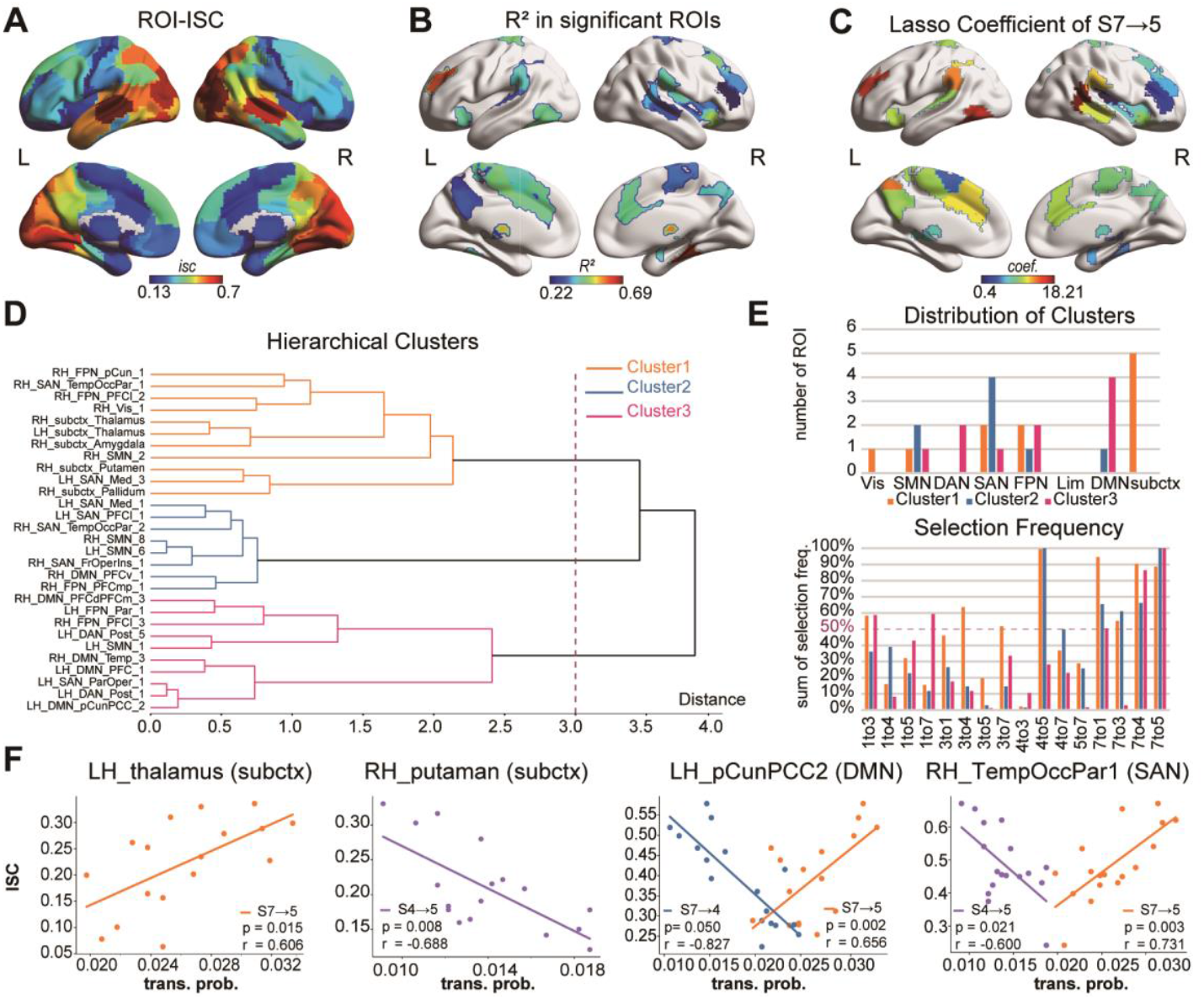
LASSO regression analysis of brain state dynamics and inter-subject correlation. (A) ROI-wise ISC analysis in the Sherlock dataset across 114 selected ROIs. (B) 29 ROIs with significant R^2^ values after FDR correction within 114 ROIs. (C) Mean coefficient distribution across ROIs, with 26 out of 29 selecting S7→5 as a feature in all 17 leave-one-out LASSO models. (D) Hierarchical clustering of the filtered coefficient matrix (ROI × valid feature, 29 × 16), revealing three clusters at a distance threshold of 3.0. (E) Additional statistics derived from LASSO analysis show that clusters primarily consist of ROIs from the SAN, DMN, and subcortex, with varying frequencies of feature selection across models. (F) Linear regression between the transition probability and the values of ISC. Four typical ROIs were selected in three clusters.

Across these 29 regions, LASSO identified a total of 16 distinct state transitions as significant predictors of ISC (Figure 3E, bottom). To further characterize the LASSO-selected brain transition patterns, we applied hierarchical clustering to the LASSO coefficient matrix (16 transitions × 29 ROIs). This clustering revealed three distinct groups (Figure 3D and 3E top): Cluster 1 primarily comprised subcortical regions, including the thalamus, amygdala, putamen and pallidus; Cluster 2 was dominated by ROIs belonging to the SAN, and Cluster 3 mainly included ROIs from the higher-order association networks, including the DMN, FPN and attention networks. The differential distribution of predictive transitions across these three clusters suggests network-specific mechanisms by which brain state dynamics influence ISC during naturalistic processing.

To examine the distribution of predictive state transitions across the three clusters, we calculated the frequency of each state transition within each cluster. Applying a 50% frequency threshold (Figure 3E bottom), we found notable differences in the number of predictive transitions across networks: Cluster 1 was characterized by 8 consistently selected transitions, while Clusters 2 and 3 showed 5 and 4 consistently selected transitions, respectively. This overrepresentation of transitions in Cluster 1 (8 vs. 5 and 4) suggests that this cluster, predominantly including the thalamus and other subcortical regions, may serve as central hubs in coordinating large-scale brain dynamics, engaging with multiple transition patterns that influence inter-subject neural synchronization. We also observed both shared and distinct state transitions across clusters. Notably, transition S7→5 showed high frequency across all three clusters, potentially representing a fundamental synchronization mechanism common to multiple brain networks. In contrast, transition S4→5 was predominantly selected in Clusters 1 and 2, while transition S7→4 was primarily selected in Clusters 1 and 3, suggesting network-specific synchronization mechanisms.

To better elucidate the relationship between selected transitions and regional ISC values, we conducted correlation analyses, as illustrated in Figure 3F, which highlights four exemplary ROIs. In Cluster 1 (subcortical regions), the S7→5 transition showed a significant positive correlation with ISC values in the left thalamus (r = 0.606, FDR-corrected P = 0.015), whereas the S4→5 transition demonstrated a significant negative correlation with ISC in the right putamen (r = -0.688, FDR-corrected P = 0.008). In Clusters 2 and 3, specific state transitions (such as S7→5 and S4→5/S7→4) exhibited significant or marginally significant linear associations with ISC values across several ROIs (see Table S1). To exemplify these associations, we highlight two representative ROIs: In LH_pCunPCC2 (encompassing the left precuneus and posterior cingulate cortex), transition S7→5 was positively correlated with ISC (r = 0.656, FDR-corrected P = 0.002), whereas S7→4 had a negative correlation (r = -0.827, FDR-corrected P = 0.050). Likewise, in RH_TempOccPar1 (primarily comprising the Temporal-Occipital-Parietal junction), S7→5 was positively correlated (r = 0.731, FDR-corrected P = 0.003), while S4→5 exhibited a negative correlation (r = -0.600, FDR-corrected P = 0.021).

### Thalamic BOLD Activity Reveals the Transition-Specific Thalamocortical Modulation

Our analysis revealed that the bilateral thalamus exhibited notably high predictive relationships with state transitions, with relatively high R^2^ values across all examined regions. This finding aligns with the thalamus’s established role as a central relay hub that integrates and routes information between cortical and subcortical structures. The thalamus’s strong involvement in our LASSO models suggests it may function as a key regulator in orchestrating transitions between different brain states, potentially controlling the flow of information that drives neural synchronization across individuals. This observation prompted us to conduct a targeted investigation into the relationship between thalamic activity and brain state dynamics. Specifically, we concentrated on the eight state transitions that exhibited over 50% selection frequency in subcortical Cluster 1, as identified by the LASSO analysis above. For each transition, we analyzed thalamic BOLD activity around state shifts to characterize how thalamic activation supports brain state transitions.

As illustrated in Figure 4, certain state transitions—including S1→3, S3→7, S7→3, and S7→5—exhibit stable significant differences in thalamic activity across the transition period. We identified two distinct thalamic activity patterns associated with these state transitions. For transitions S1→3 and S3→7, thalamic activity showed a marked increase prior to the transition, with significant differences emerging approximately 3–4 TRs before the transition point. This suggests a preparatory role for thalamic regions in facilitating these state changes. In contrast, during transitions such as S7→3 and S7→5, thalamic activity is significantly suppressed before the transition, followed by a rapid post-transition increase that surpasses baseline levels, which may reflect a disengagement from the preceding state and subsequent engagement in the new state, thus supporting functional flexibility during dynamic state switching. For the two state transitions, S1 → 3 and S7 → 5, that showed the most pronounced differences, we performed further analyses and confirmed consistent changes in thalamic activity across both hemispheres and among different thalamic cell populations (core and matrix cells, Figure S2A). These results highlight a strong thalamic involvement in shaping the overall dynamics of brain state changes.

**Figure 4.**
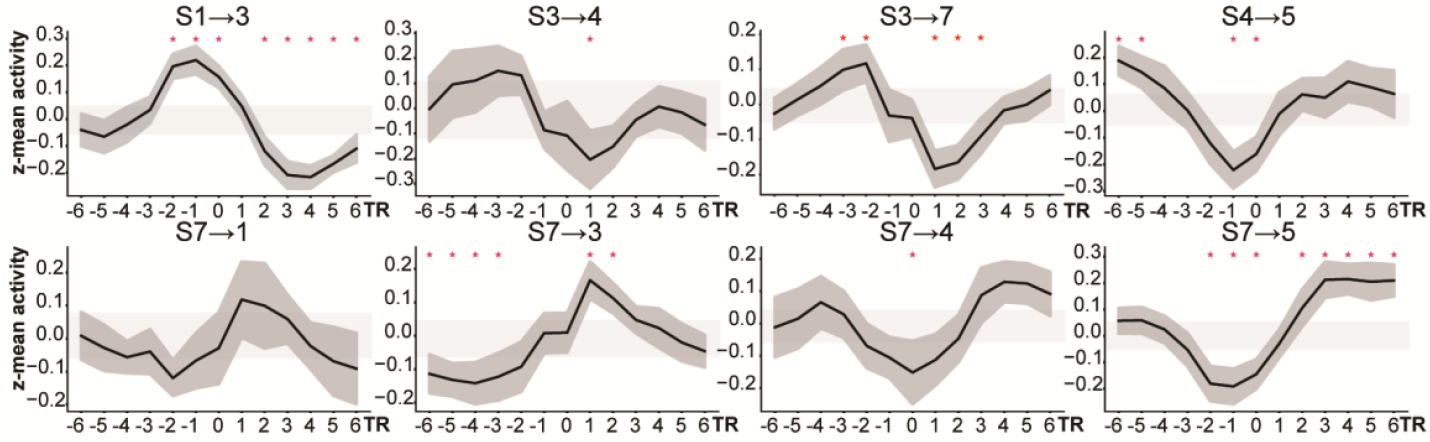
Brain state transitions are accompanied by changes in thalamic activity. Thalamic BOLD activity associated with eight state transitions was assessed across a 13-TR window centered on the transition onset (0 TR). The solid black line depicts the average BOLD response across participants, while the shaded gray band denotes the 95% confidence interval. The light gray region marks the upper and lower limits established through permutation testing. At each time point, independent t-tests compared the data from 17 subjects with 1,000 permutations; red stars indicate time points with significant differences (p < 0.001, Bonferroni-corrected for 13 comparisons).

**Figure 5.**
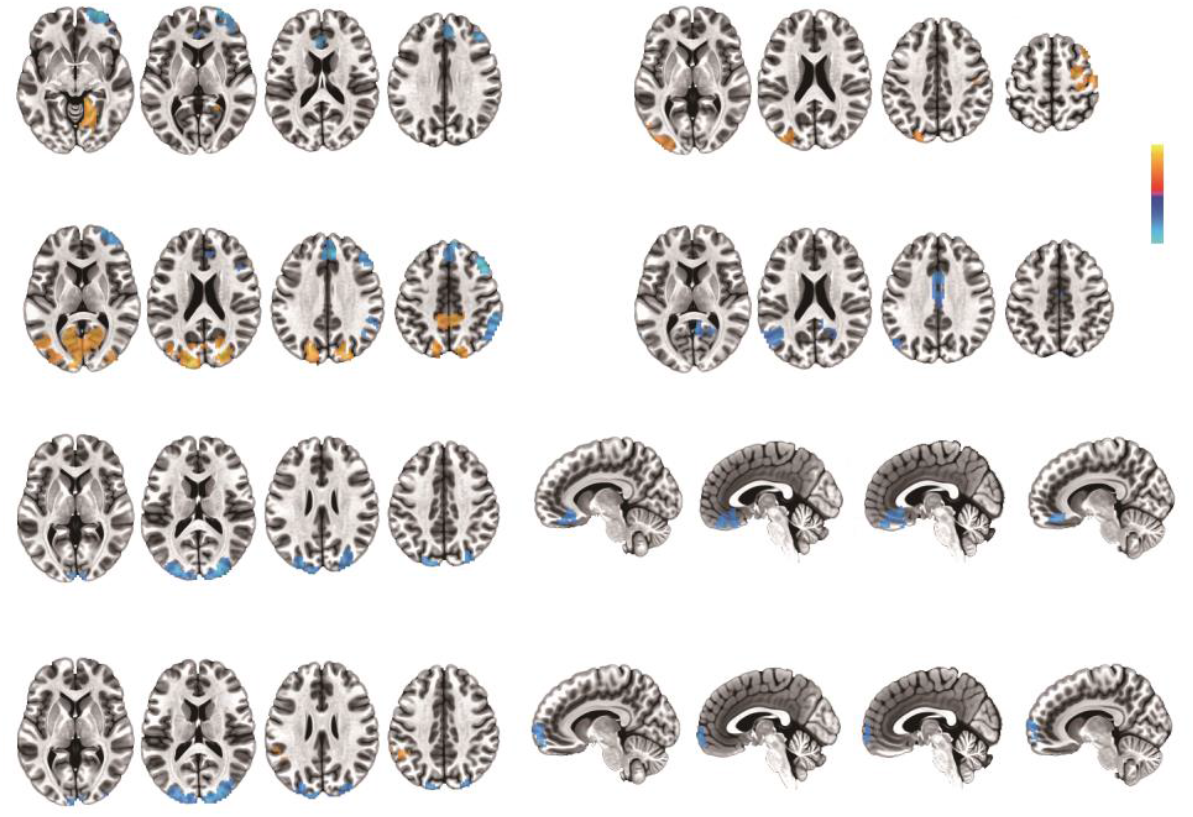
Thalamocortical functional connectivity changes at brain state transitions. Thalamocortical functional connectivity at brain state transitions was compared against null distributions generated by randomly placed transitions. Voxels with |*z*| > 1.95 (*ps*. < 0.025) are highlighted, with clusters corrected using AFNI (NN1, bi-side, 201 voxels). For S7→5, a more lenient threshold of |*z*| > 1.645 was applied to detect marginally significant clusters (*ps*. < 0.05) as we did not find any valid cluster at the original threshold.

### Thalamocortical Functional Connectivity Changes at Brain State Transitions

Next, we investigated the changes in functional connectivity patterns between the thalamus and cortical regions during brain state transitions. Given that thalamic subregions containing PVALB-enriched core cells and CALB1-enriched matrix cells are known to play distinct roles in regulating cortical activity, we extracted the average signals from these subregions to characterize their distinct connectivity profiles around state transition points. Although the activity levels of these two subregions were not markedly different near transitions, as shown in Figure S2A, we observed significant differences in their functional connectivity at these transition points. For clarity, we concentrated our analysis on two representative state transitions with contrasting thalamic activity signatures: S1→3 and S7→5. During S1→3 transitions, both thalamic cell populations exhibited comparable connectivity alterations at state onset, including increased FC with primary visual areas and decreased FC with anterior/superior frontal regions. Notably, matrix cells demonstrated additional connectivity changes, such as enhanced FC with the occipital cortex and precuneus and reduced FC with the right inferior parietal lobule. In contrast, S7→5 transitions revealed a different pattern: core cells showed heightened connectivity with visual and precentral regions, while matrix cells displayed reduced connectivity with the left inferior parietal lobule and posterior cingulate cortex. These findings underscore the distinct roles of thalamic cell populations in cortical network reconfiguration during specific brain state transitions. Additionally, we compared the transitions S4→5 and S7→4 and found no significant differences in the functional connectivity of core and matrix cells during these transitions.

## Discussion

Applying pairwise maximum entropy modeling on the Sherlock dataset, we identified two dominant brain states: a perceptually and attentionally engaged state (SAN/DAN/SMN/Vis) and a state dominated by introspective and integrative processing (DMN/Lim/FPN). First, route-specific BOLD modulation was observed within canonical brain systems: “Easy” routes primarily recruited DMN hubs (such as the precuneus and MTG), while “Hard” routes engaged regions within the multiple-demand network (such as IFG and IPL). Second, we found that the relationship between state transition probability and regional ISC was significantly identified in 29 ROIs, which could be further classified into three groups (subcortical-rich, SAN-rich, DMN-rich). Third, high consistency in cross-subject thalamic BOLD activity changes was observed near these transitions that predict regional ISC, especially at S7→S5 and S1→S3, revealing thalamus-guided thalamocortical modulations. Finally, thalamocortical dynamic functional connectivity between core/matrix cells and cortical areas showed distinct connectivity patterns at transitions S1→S3 and S7→S5: core cells modulated primary visual areas while matrix cells modulated DMN regions. Overall, these findings demonstrate that (1) cortical state transitions are orchestrated by specialized neural substrates; (2) intersubject synchronization dynamically relates to transition probabilities; and (3) thalamocortical circuits facilitate state transitions via temporally precise, nucleus-specific interactions.

Our energy landscape analysis identified two dominant, anti-correlated brain states that reflect fundamental processing modes during naturalistic viewing: State 1 (involving the salience, dorsal attention, somatomotor, and visual networks; SAN/DAN/SMN/Vis), and State 7 (engaging the default mode, limbic, and frontoparietal networks; DMN/LIM/FPN). These findings align with previous resting-state studies that utilized the pair-wise maximum entropy model (Wu et al., 2025; Yamashita et al., 2021), suggesting that the identified brain states remain consistent when subjects are engaged in naturalistic movie viewing. This functional dichotomy aligns with the ‘dual intertwined architecture’ hypothesis (Mesmoudi et al., 2013; Yamashita et al., 2021). In this framework, State 7 may support multitemporal integration—the combination of information across different time scales—while State 1 may facilitate multimodal integration, meaning the synthesis of input from various sensory modalities. Beyond identifying these states, we observed distinctive features in their transition dynamics. First, transitions between the dominant states exhibited a clear hierarchical pattern: direct State 1↔State 7 transitions were significantly less frequent than transitions that passed through intermediate states (State 3↔State 5; Figure 1C). This staged transition pattern suggests that the brain minimizes energy barriers by traversing hybrid network configurations (Deco and Kringelbach, 2016), which functionally enable a flexible shifting between external sensory engagement and internal cognitive integration critical for naturalistic viewing. These intermediate states may serve as functional bridges that coordinate the temporal sequencing and sensory modalities required by complex audiovisual stimuli. Second, the intermediate states (States 3, 4, and 5), which together account for approximately 45% of viewing time, exemplify a trajectory consistent with contemporary models of brain dynamics as metastable. Rather than shifting abruptly between dominant states, the brain transitions smoothly through these intermediate configurations, reflecting the fluid and flexible nature of neural state changes (Saggar et al., 2022). Such metastability may be especially adaptive for processing audiovisual narratives, which require the brain to maintain both external focus and internal interpretation (Deco and Jirsa, 2012), highlighting the functional importance of these intermediate states as facilitators for integrating sensory inputs with higher-order cognitive processes during natural viewing.

The classification of energy landscape transitions into “Hard” (energy-increasing) and “Easy” (energy-decreasing) routes reveals a fundamental duality in cognitive state switching, whereby these pathways engage distinct neural systems with complementary functional roles. Specifically, easy routes (energy minimization) preferentially activated the DMN regions, particularly the precuneus and middle temporal gyrus. These “easy” transition routes likely require minimal cognitive control and attentional resource allocation. In these contexts, the DMN, which has been recognized as task-negative and associated with internally focused thought (Buckner et al., 2008; Raichle et al., 2001), is permitted to remain active, thereby creating an opportunity for brief, low-effort introspective processing that helps participants assimilate plot developments, update their personal understanding of the narrative, and maintain psychological continuity. This finding is consistent with the recently established role of the DMN in narrative maintenance and interpreting complex (Lee and Chen, 2024; Su et al., 2025), temporally extended natural experiences. In contrast, during “hard” (energy-increasing) state transitions, presumably when complex scene changes, surprise events, or heightened attentional demands occur, we found that instead of DMN activations, the multiple demand network (MDN) regions, including the inferior frontal gyrus (IFG) and inferior parietal lobule (IPL), are robustly recruited, showing significant increases in activation immediately after transitions. This complementarity aligns with principles of dynamical systems, where easy transitions reflect a “gradient descent” into stable attractor states, whereas hard transitions indicate points that require energetic input to overcome network barriers. More broadly, this complementary recruitment of DMN and MDN under naturalistic conditions illustrates how large-scale brain dynamics are flexibly organized to balance internal situation modeling with responsiveness to environmental change, a principle likely essential for navigating real-world cognitive challenges.

The LASSO regression revealed a systematic relationship between cortical state transition dynamics and intersubject correlation patterns across three distinct neural systems (subcortical-rich, SAN-rich, DMN-rich). Remarkably, the ISC of subcortical regions could be reliably predicted by these transition probabilities, suggesting that subcortical structures actively participate in, and perhaps even facilitate, large-scale state transitions (Aguilar and McNally, 2022). Importantly, these findings emerged despite defining states solely by cortical features, highlighting the often-underestimated role of subcortical regions in orchestrating brain-wide dynamics (Favaretto et al., 2022). Furthermore, specific transitions such as S7→S5, S7→S4 and S4→S5 showed robust predictive power for regional ISC. Of particular note, the S7→S5 transition was associated with enhanced neural synchronization across participants, whereas transitions S7→S4 and S4→S5 were linked to reduced intersubject synchrony (negative coefficients). From the perspective of large-scale brain networks, each state transition likely involves distinct patterns of engagement and disengagement across functional systems. For example, the S7→S5 transition may indicate a shift from an introspective, integrative state, which is often associated with the DMN, situated at the highest hierarchical level (Baldassano et al., 2017; Geerligs et al., 2022; Margulies et al., 2016), toward an attention-driven state supported by the central executive and attention networks. This reconfiguration may enable a more efficient and focused response to salient external stimuli, facilitating synchronized cognitive engagement and underpinning moments of collective focus or shared understanding during narrative experiences. Conversely, transitions like S7→S4 and S4→S5 might involve partial decoupling from the current state to a new one, or the recruitment of more flexible, intermediate networks such as the salience or frontoparietal control networks. During these transitions, attentional engagement may become more variable, allowing individuals to drift into personal reflection, subjective interpretation, or fluctuating focus. As a result, intersubject neural synchronization decreases, highlighting increased individuality in how each person engages with the narrative. These nuanced dynamics underscore the complex interplay between stable network configurations and flexible transitions that enable both shared understanding and personalized interpretation within social and cognitive contexts. Further research should explore the mechanistic underpinnings of these transitions and their behavioral correlates.

Our thalamic analyses yielded two key insights into the neural dynamics of state transitions and provided new evidence for thalamocortical modulation. First, we identified two main BOLD activity patterns near the 8 selected state transitions. The first pattern showed increased/peaked activity before transition onset (e.g., S1→S3), while the second pattern exhibited increased/peaked activity after transition onset (e.g., S7→S5). These two distinct patterns suggest the thalamus may actively initiate or coordinate these shifts, rather than simply following cortical changes. Second, we observed cell population-specific modulation effects in thalamocortical functional connectivity patterns during brain state transitions. Matrix cell-specific thalamic subregions showed transition-dependent coupling with the posterior medial cortex, which serves as the posterior hub region of the DMN . For example, there was increased functional connectivity with the precuneus during S1→S3, and decreased connectivity with the posterior cingulate during S7→S5. In contrast, core cell-specific thalamic subregions increased connectivity with the primary sensory regions, reflecting their distinct functional roles (Halassa and Kastner, 2017; Hwang et al., 2017; Müller et al., 2020; Raut et al., 2020; Saalmann et al., 2012). These findings align with and refine the thalamocortical framework (Shine, 2021), which conceptualizes the thalamus as a central regulator of large-scale brain state transitions through flexible integration and segregation of cortical networks. Our findings extend this framework by showing that the thalamus plays an active role in modulating cortical dynamics during naturalistic stimuli, with matrix- and core-specific thalamic subcircuits preferentially routing information to distinct brain networks, thereby supporting different brain state transitions.

Our study has several limitations. First, while Lasso regression was applied to identify state transitions potentially related to regional ISC, several limitations should be acknowledged. ISC values in subcortical regions may be less reliable compared to those in early sensory and higher-order DMN regions due to lower signal-to-noise ratios and susceptibility to physiological artifacts. Therefore, we interpret the relationship between state transition probability and regional ISC as exploratory rather than treating transition probability as a definitive predictor of ISC. Additionally, our analysis reveals associations rather than causal relationships, and the temporal resolution of fMRI may not capture the full dynamics of rapid neural transitions. Future studies should consider these methodological constraints when interpreting thalamocortical state transition mechanisms. Second, extracting subcortical signals, especially specific small nuclei, from low-resolution 3Tesla fMRI data remains challenging. Although this study analyzed signals at the ROI or thalamic population (matrix and core) level with brain imaging resolution of 3 mm^3^, and previous studies have validated the feasibility of using similar resolution (Tu et al., 2024; Wen et al., 2023), finer-grained investigations focusing on specific thalamic nuclei are still warranted. Prior research using 7 Tesla fMRI data indicates thalamocortical projections may be nucleus-specific(Setzer et al., 2022). Further research could build on our findings to further elucidate the relationship between cognitive functions and thalamocortical modulation.

## Acknowledgement

We would like to thank Dr.Chen, Dr.Ezaki and Dr.Watanabe for sharing data and providing valuable suggestions on methods. This work was supported by the National Natural Science Foundation of China (https://www.nsfc.gov.cn/, grant numbers 82072000), the Natural Science Foundation of Heilongjiang Province, China (http://kjt.hlj.gov.cn/, grant number YQ2023H016), the Fundamental Research Funds for the Central Universities (Grant No. HIT.OCEF.202301).

