## Supplementary Materials for "Energy Landscape Analysis Reveals Thalamic Modulation of Brain State Transitions During Movie Watching"

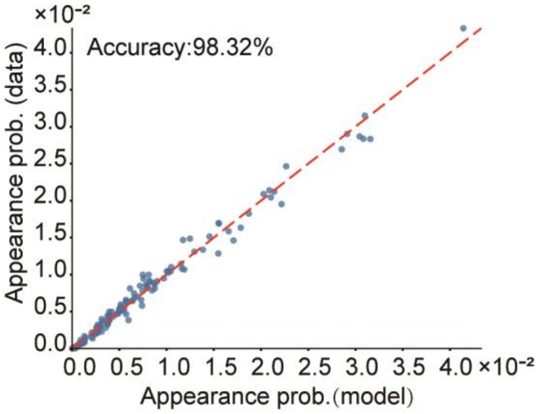


**Figure S1 The pairwise maximum entropy model fitted well with empirical data.**

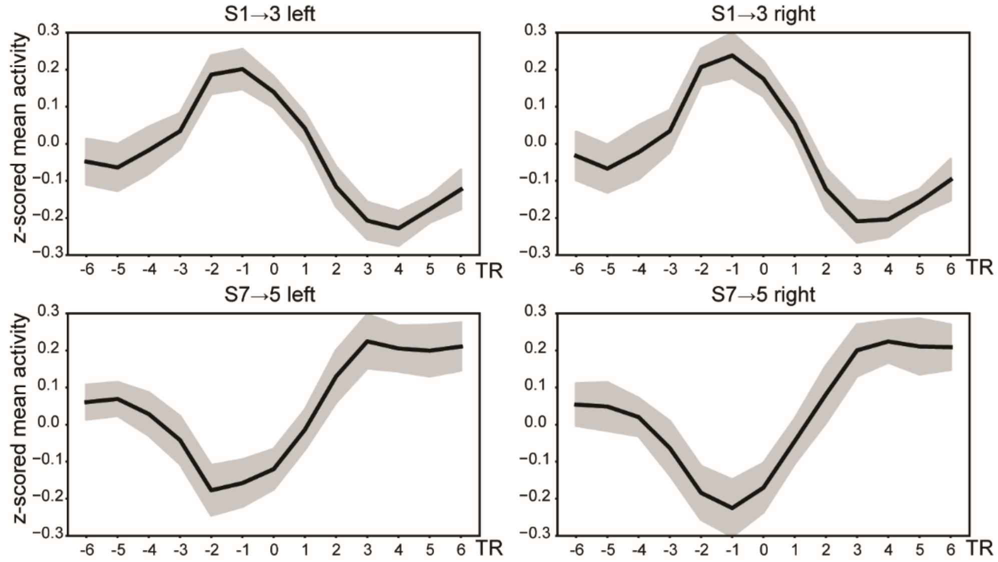


**Figure S2 Transition-triggered BOLD activity in bilateral thalamus.**

Thalamic BOLD activity across a 13-TR window centered on new state onset. The solid black line represents the mean z-scored BOLD activity across subjects, with the dim gray band indicating the 95% confidence interval.

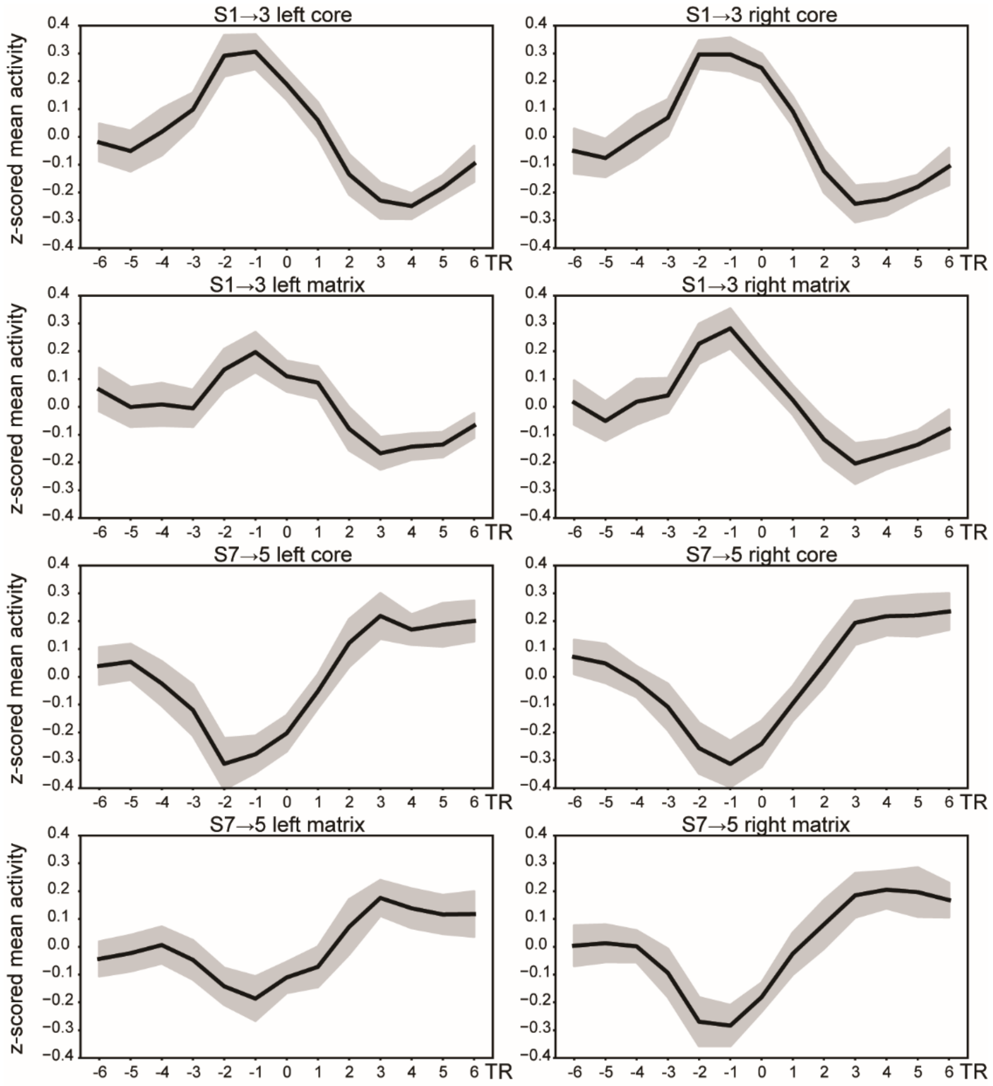


**Figure S3 Transition-triggered BOLD activity of the two main cell populations in the bilateral thalamus.**
The activity was extracted with an open-sourced CALB1 minus PVALB thalamic atlas (Müller et al., 2020), where the core cell was identified by a negative value while the matrix cell population was identified by a positive value. Thalamic BOLD activity across a 13-TR window centered on new state onset. The solid black line represents the mean z-scored BOLD activity across subjects, with the dim gray band indicating the 95% confidence interval.

**Table S1 Significances of correlation between transition probability and ISC values in filtered out ROIs**

| ROI | 1to3 | 1to4 | 1to5 | 1to7 | 3to1 | 3to4 | 3to5 | 3to7 | 4to3 | 4to5 | 4to7 | 5to7 | 7to1 | 7to3 | 7to4 | 7to5 |
|---|---|---|---|---|---|---|---|---|---|---|---|---|---|---|---|---|
| LH_SMN_1 | 0.907 | 0.798 | 0.881 | 0.953 | 0.640 | 0.957 | 0.959 | 0.909 | 0.883 | 0.079 | 0.907 | 0.557 | 0.987 | 0.843 | 0.050 | 0.036 |
| LH_SMN_6 | 0.825 | 0.798 | 0.881 | 0.953 | 0.485 | 0.957 | 0.959 | 0.909 | 0.864 | 0.009 | 0.974 | 0.515 | 0.987 | 0.843 | 0.164 | 0.007 |
| LH_DAN_Post_1 | 0.825 | 0.932 | 0.881 | 0.953 | 0.485 | 0.957 | 0.975 | 0.909 | 0.883 | 0.055 | 0.907 | 0.515 | 0.987 | 0.843 | 0.050 | 0.002 |
| LH_DAN_Post_5 | 0.825 | 0.932 | 0.881 | 0.953 | 0.485 | 0.957 | 0.975 | 0.909 | 0.883 | 0.297 | 0.954 | 0.515 | 0.987 | 0.874 | 0.121 | 0.008 |
| LH_SAN_ParOper_1 | 0.825 | 0.871 | 0.881 | 0.953 | 0.485 | 0.957 | 0.975 | 0.909 | 0.883 | 0.077 | 0.907 | 0.515 | 0.995 | 0.843 | 0.071 | 0.002 |
| LH_SAN_PFCl_1 | 0.825 | 0.801 | 0.928 | 0.953 | 0.485 | 0.957 | 0.975 | 0.909 | 0.864 | 0.011 | 0.907 | 0.515 | 0.987 | 0.874 | 0.064 | 0.002 |
| LH_SAN_Med_1 | 0.825 | 0.798 | 0.881 | 0.953 | 0.566 | 0.957 | 0.959 | 0.909 | 0.864 | 0.008 | 0.907 | 0.705 | 0.987 | 0.874 | 0.050 | 0.014 |
| LH_SAN_Med_3 | 0.907 | 0.871 | 0.881 | 0.953 | 0.943 | 0.957 | 0.975 | 0.909 | 0.884 | 0.008 | 0.907 | 0.668 | 0.987 | 0.843 | 0.050 | 0.050 |
| LH_FPN_Par_1 | 0.825 | 0.917 | 0.881 | 0.953 | 0.485 | 0.957 | 0.975 | 0.909 | 0.883 | 0.174 | 0.907 | 0.557 | 0.987 | 0.843 | 0.064 | 0.007 |
| LH_DMN_PFC_1 | 0.825 | 0.874 | 0.995 | 0.953 | 0.485 | 0.957 | 0.975 | 0.909 | 0.864 | 0.058 | 0.907 | 0.557 | 0.987 | 0.880 | 0.050 | 0.002 |
| LH_DMN_pCunPCC_2 | 0.825 | 0.932 | 0.881 | 0.953 | 0.609 | 0.957 | 0.975 | 0.909 | 0.883 | 0.095 | 0.907 | 0.515 | 0.987 | 0.843 | 0.050 | 0.002 |
| RH_Vis_1 | 0.907 | 0.871 | 0.881 | 0.953 | 0.530 | 0.957 | 0.975 | 0.909 | 0.864 | 0.008 | 0.907 | 0.557 | 0.987 | 0.843 | 0.001 | 0.008 |
| RH_SMN_2 | 0.905 | 0.798 | 0.881 | 0.953 | 0.485 | 0.957 | 0.959 | 0.909 | 0.864 | 0.009 | 0.907 | 0.791 | 0.987 | 0.843 | 0.043 | 0.036 |
| RH_SMN_8 | 0.825 | 0.798 | 0.881 | 0.953 | 0.558 | 0.957 | 0.959 | 0.909 | 0.864 | 0.009 | 0.907 | 0.515 | 0.987 | 0.874 | 0.121 | 0.008 |
| RH_SAN_TempOccPar_1 | 0.825 | 0.801 | 0.881 | 0.953 | 0.485 | 0.957 | 0.959 | 0.909 | 0.864 | 0.021 | 0.907 | 0.557 | 0.987 | 0.843 | 0.050 | 0.003 |
| RH_SAN_TempOccPar_2 | 0.825 | 0.801 | 0.881 | 0.953 | 0.485 | 0.957 | 0.959 | 0.909 | 0.864 | 0.010 | 0.907 | 0.515 | 0.987 | 0.843 | 0.050 | 0.002 |
| RH_SAN_FrOperIns_1 | 0.825 | 0.801 | 0.881 | 0.953 | 0.485 | 0.957 | 0.975 | 0.909 | 0.864 | 0.008 | 0.974 | 0.557 | 0.987 | 0.843 | 0.050 | 0.007 |
| RH_FPN_PFCl_2 | 0.825 | 0.798 | 0.881 | 0.953 | 0.485 | 0.957 | 0.975 | 0.909 | 0.864 | 0.008 | 0.907 | 0.754 | 0.987 | 0.843 | 0.043 | 0.024 |
| RH_FPN_PFCl_3 | 0.825 | 0.798 | 0.881 | 0.953 | 0.485 | 0.957 | 0.975 | 0.909 | 0.864 | 0.058 | 0.907 | 0.592 | 0.987 | 0.843 | 0.043 | 0.017 |
| RH_FPN_PFCmp_1 | 0.825 | 0.801 | 0.881 | 0.953 | 0.485 | 0.957 | 0.959 | 0.909 | 0.864 | 0.008 | 0.907 | 0.705 | 0.987 | 0.843 | 0.050 | 0.014 |
| RH_FPN_pCun_1 | 0.825 | 0.844 | 0.881 | 0.953 | 0.485 | 0.957 | 0.975 | 0.909 | 0.864 | 0.027 | 0.907 | 0.515 | 0.987 | 0.843 | 0.043 | 0.005 |
| RH_DMN_Temp_3 | 0.825 | 0.871 | 0.881 | 0.953 | 0.378 | 0.957 | 0.959 | 0.994 | 0.883 | 0.175 | 0.907 | 0.515 | 0.987 | 0.843 | 0.313 | 0.002 |
| RH_DMN_PFCv_1 | 0.907 | 0.798 | 0.881 | 0.953 | 0.640 | 0.957 | 0.959 | 0.909 | 0.864 | 0.008 | 0.907 | 0.664 | 0.995 | 0.874 | 0.043 | 0.008 |
| RH_DMN_PFCdPFCm_3 | 0.825 | 0.871 | 0.881 | 0.953 | 0.499 | 0.957 | 0.975 | 0.909 | 0.864 | 0.192 | 0.907 | 0.557 | 0.987 | 0.843 | 0.050 | 0.015 |
| LH_Sub_Thalamus | 0.825 | 0.871 | 0.881 | 0.953 | 0.806 | 0.957 | 0.975 | 0.909 | 0.864 | 0.307 | 0.954 | 0.515 | 0.987 | 0.874 | 0.164 | 0.015 |
| RH_Sub_Thalamus | 0.825 | 0.871 | 0.881 | 0.953 | 0.732 | 0.957 | 0.959 | 0.909 | 0.864 | 0.140 | 0.907 | 0.557 | 0.987 | 0.843 | 0.121 | 0.016 |
| RH_Sub_Putamen | 0.900 | 0.871 | 0.881 | 0.953 | 0.838 | 0.957 | 0.959 | 0.909 | 0.864 | 0.008 | 0.954 | 0.557 | 0.987 | 0.843 | 0.121 | 0.031 |
| RH_Sub_Pallidum | 0.907 | 0.798 | 0.881 | 0.953 | 0.970 | 0.957 | 0.959 | 0.909 | 0.864 | 0.008 | 0.907 | 0.756 | 0.987 | 0.843 | 0.124 | 0.098 |
| RH_Sub_Amygdala | 0.825 | 0.871 | 0.995 | 0.953 | 0.908 | 0.997 | 0.959 | 0.909 | 0.883 | 0.116 | 0.907 | 0.515 | 0.987 | 0.843 | 0.121 | 0.027 |
